# Diastolic Dysfunction Accompanies Alterations in Myocardial Structure, Cellular Composition and Macrophage Polarization in Survivors of Ionizing Radiation Exposure

**DOI:** 10.1101/2020.02.18.953190

**Authors:** Kristofer T. Michalson, Gregory O. Dugan, David L. Caudell, J. Mark Cline, Dalane W. Kitzman, Thomas C. Register

**Author notes:** Address for correspondence: Department of Pathology, Section on Comparative Medicine, Wake Forest University School of Medicine, Medical Center Boulevard, Winston-Salem, NC 27157-1040;.

## Abstract

**Rationale:** Radiation induced heart disease (RIHD) is a significant delayed/late effect of ionizing radiation exposure.

**Objective:** To determine the cardiac effects of total body irradiation (TBI) in male rhesus macaques, a translational non-human primate (NHP) model.

**Methods and Results:** Echocardiography was performed on survivors of a single dose (6.4-8.5 Gy) of TBI (n=34) and non-irradiated controls (n=26) divided into longer (LT IRR) and shorter term (ST IRR) survivors and controls to assess the effects of time since TBI on phenotypes. LT IRR had increased Doppler transmitral early filling velocities (E), decreased early mitral annular descent velocities (e’), and higher E/e’ ratio compared to LT CTL (all p≤0.05), indicating left ventricular (LV) diastolic dysfunction. Echocardiographic stroke volume, cardiac output, and end-diastolic and systolic volumes were also lower in LT IRR than controls (all p ≤ 0.05). ST IRR had similar alterations in LV diastolic function but not in cardiac volumetric measures. Analyses of LV, interventricular septum (IVS), and right ventricle (RV) myocardium from deceased irradiated animals (n=17) exposed to a single dose (6.9-8.05 Gy) TBI and non-irradiated controls (n=12) showed that IRR animals had decreased LV and IVS capillary density, and increased LV fibrosis, pan-cardiac fibroblast and macrophage staining, LV and IVS M2 macrophages, and pan-cardiac M1 macrophages (all p<0.05). While M2 predominated over M1 macrophages in both groups, M1 showed greater increases than M2 in IRR.

**Conclusions:** LV diastolic dysfunction due to radiation exposure may be due to a combination of capillary rarefication, activation and expansion of fibroblasts, and increased accumulation of both pro-fibrotic and pro-inflammatory macrophages, all of which lead to increased myocardial stiffness due to fibrosis. Collectively, these findings provide insights into the delayed effects of acute ionizing radiation exposure and suggest that therapies targeting macrophage regulation of fibrosis may mitigate radiation induced heart disease.

## INTRODUCTION

Radiation-induced heart disease (RIHD) can be a delayed effect of acute ionizing radiation exposure (DEARE) of the heart, which can occur through accidental or malicious nuclear events, radiotherapies for lung or breast cancers and Hodgkin’s disease, or during space flight. The condition is characterized by a broad spectrum of pathologies including pericarditis (primarily at total doses greater than 40 Gy)^1^, arrhythmias, valvular disease, precocious coronary artery atherosclerosis, and myocardial fibrosis in humans.^2-4^ In humans, the delayed effects of acute radiation exposure (DEAREs) are perhaps best documented in the Life Span Study (LSS) of the Japanese survivors of the Hiroshima and Nagasaki atomic bombings, who generally received an estimated ≤2 Gy total body irradiation.^8^ Studies of the LSS showed markedly increased predispositions for heart and cardiovascular disease ^9^ as well as cancer^10^ and noncancer diseases including stroke,^11, 12^ chronic kidney disease,^13, 14^ and respiratory disease^15^.

Despite recognition of RIHD as a significant clinical concern, there are no consensus recommendations on prevention or treatment of RIHD, due in large part to significant gaps in knowledge regarding the initiation and progression of local and systemic processes involved in RIHD, particularly due to lack of animal models truly relevant to humans. These gaps include the pathogenesis of myocardial remodeling, the effects of RIHD on systolic and diastolic function, whether these changes result in heart failure (HF) with reduced ejection fraction (HFrEF) or heart failure with preserved ejection fraction (HFpEF), and how the condition progresses over time.

Myocardial fibrosis is a key pathologic phenotype of RIHD leading to adverse myocardial remodeling, but the mechanisms underlying its development, particularly the critical events involved and pertinent signaling pathways which might be targeted therapeutically are not fully understood, and initiating signals and cells promoting fibrosis are largely unknown. Our laboratory recently demonstrated that the key protein responsible for monocyte chemotaxis (MCP1) was elevated (p<0.05) in previously irradiated male rhesus macaques years after irradiation (5.6-9.7 years), suggesting a long term pro-inflammatory systemic environment in which monocyte/macrophages may play a role in the pathogenesis of RIHD fibrosis.^23^ We also recently discovered there are alterations in circulating monocyte polarization from classical towards intermediate monocyte phenotypes after 6 months after total body irradiation, providing a potential mechanism for RIHD fibrosis in the heart and other organs^25^.

The goal of this study was to determine the effects of TBI on 1) the structure and function of the heart using echocardiography and 2) the cellular and extracellular matrix composition of the myocardium using immunohistochemistry and histology to determine alterations which might play a role in RIHD. This study utilized a unique resource of male rhesus macaque survivors, termed the Radiation Survivor Cohort (RSC), that were previously exposed to single dose total body irradiation (TBI).^23, 30-32^ For functional outcomes, we specifically focused on two subsets of the RSC which differed in time elapsed since irradiation to determine the potential for long term progression which might lead to heart failure.

## METHODS

### Animals

Subjects were male rhesus nonhuman primates (NHP; *Macaca mulatta*) obtained from the Armed Forces Radiobiology Research Institute (AFRRI), the University of Maryland (UMD), the University of Illinois at Chicago (UIC), the Lovelace Respiratory Research Institute, and the Michale E. Keeling Center for Comparative Medicine and Research. Animals were screened by intradermal tuberculin testing three times a year to exclude tuberculosis, and by immunoassay upon arrival for simian retroviruses (Washington National Primate Research Center, Seattle, WA). Animals were aged based on dentition, weighed, examined by a veterinarian and quarantined for 60 days upon arrival per standard institutional protocol. The animals were assessed and treated for discomfort and illness as described previously by DeBo^23^ including criteria for humane euthanasia. All experimental procedures were conducted in compliance with the Wake Forest School of Medicine (WFSM) Institutional Animal Care and Use Committee requirements and followed the Guide for Care and Use of Laboratory Animals.

### In vivo: Echocardiographic phenotyping

Two groups of animals were chosen from our available dataset of 60 NHP for the current study of echocardiographic phenotypes, 1) an older, longer term survivor group of 14 mature adult (12.6 to 20.3 year old) long-term survivors (LT IRR) exposed to a single TBI dose of 6.5 to 8.4 Gy approximately 9.7 to 11.6 years earlier, which were compared to 15 age-matched, nonirradiated controls (LT CTL)(Table 1), and younger, shorter term survivor group (ST IRR) of 20 young adult (4.7 to 6.0 year old) exposed to a single TBI dose of 6.4-8.5 Gy approximately 2.16 to 2.96 years earlier, which were compared to 10 age-matched, nonirradiated controls (ST CTL)(Table 1).

**Table 1:** Animal Demographics of Previously Irradiated Long Term, Short Term, and Deceased Male Rhesus Macaque Survivors (LT IRR, ST IRR, Deceased IRR) and Controls (LT CTL, ST CTL, Deceased CTL)

|  | LT IRR<br>(N=14) | LT CTL<br>(N=15) | P | ST IRR<br>(N=20) | ST CTL<br>(N=10) | P | Deceased<br>IRR (N=17) | Deceased<br>CTL (N=12) | P |
| --- | --- | --- | --- | --- | --- | --- | --- | --- | --- |
| Age (Years) | 14.96 (0.63) | 14.15 (0.28) | 0.25 | 5.47 (0.08) | 5.25 (0.33) | 0.40 | 12.14 (0.96) | 9.33 (0.92) | 0.47 |
| Time Since<br>Irradiation<br>(Years) | 10.48 (0.18) | --- | --- | 2.51 (0.07) | --- | --- | 5.80 (0.55) | --- | --- |
| Dose (Gy) | 7.40 (0.16) | --- | --- | 7.01 (0.14) | --- | --- | 6.93 (0.13) | --- | --- |
| Body Weight<br>(kg) | 11.31 (1.11) | 14.62 (1.25) | 0.06 | 9.22 (0.56) | 8.77 (0.35) | 0.86 | 13.25 (0.85) | 10.90 (0.54) | **0.04<sup>a</sup>** |
| Body Surface<br>Area (m<sup>2</sup>) | 0.48 (0.03) | 0.55 (0.03) | 0.16 | 0.42 (0.29) | 0.41 (0.01) | 0.97 | 0.54 (0.02) | 0.47 (0.02) | 0.06 |

Data presented as mean (SEM)

### Ex vivo: Histochemical and immunohistochemical phenotyping of the heart

Cardiac tissues were obtained from a separate set of animals that had died or been euthanized due to morbidity. All subjects underwent a complete necropsy with saline flush as detailed in DeBo et al.^23^ Subjects included 17 irradiated animals (Deceased IRR) exposed to a single TBI dose of 6.92 to 8.05 Gy TBI approximately 2.1 to 10.6 years prior that died or were euthanized at 5.6 to 19.2 years of age (mean 12.1 years), along with 12 non-irradiated controls (Deceased CTL) that died or were euthanized at 6.8 to 17.3 years of age (mean 9.3 years)(Table 1). Longitudinal sections of LV, interventricular septum (IVS) and right ventricle (RV) were sectioned from paraformaldehyde fixed hearts as previously described.^40^

### Diet

Prior to arrival at WFSM (and during the time of irradiation), animals consumed standard laboratory primate chow diet (18% protein, 13% fat and 69% carbohydrates, with a low cholesterol content of 0.023 mg/Cal (for example, Monkey Diet 5038; LabDiet®, St. Louis, MO). After arrival at WFSOM, monkeys were transitioned to a moderately atherogenic Western diet developed by WFSM and Purina® (Typical American Diet, TAD; Purina LabDiet as NHP Diet) which contained 18.4% protein, 36.2% fat and 45.4% carbohydrates as percentage of calories provided, and a cholesterol content of 0.18 mg/Cal. Any animals with a hemoglobin A1c higher than 6.5%, indicating diabetes mellitus onset, were transitioned to standard laboratory chow in the interest of health preservation.

### In Vivo: Echocardiography

Animals were sedated with 10-15 mg/kg ketamine and 0.05-0.15 mg/kg midazolam to facilitate echocardiography using a Logiq S8 ultrasound system (GE Healthcare, Chicago, ILL) by trained NHP ultrasonographers following guidelines for image acquisition and analysis established by the American Society of Echocardiography and European Association of Cardiovascular Imaging^41, 42^ combined with recommendations specifically for rhesus macaque echocardiograms reported by Korcarz et al.^43^ Captured images and video recordings were subsequently analyzed using Image-Arena software (TomTec Imaging Systems GMBH, Munich, Germany). Heart rate and blood pressure was obtained throughout the echocardiogram using electrocardiography and blood pressure cuff respectively. Parasternal short and long axis, and apical two (A2C) and four chamber (A4C) views of the heart were obtained to evaluate structure and systolic and diastolic function. Parasternal short axis brightness (B) mode views at the level of the aortic valve were used to measure end systolic aortic root and left atrial diameters to calculate the aortic root (AO) to left atrium (LA) diameter ratio (AO/LA). Motion (M) mode views at the level of the LV papillary muscles were captured to assess interventricular septal thicknesses at end diastole (IVSd) and end systole (IVSs), LV internal diameters at end diastole (LVIDd) and end systole (LVIDs), LV posterior wall thicknesses at end diastole (LVPWd) and end systole (LVPWs), and fractional shortening (FS). The LV endocardium was traced at end diastole and end systole in A2C and A4C views to obtain end diastolic (EDV) and systolic volumes (ESV) by method of disks, stroke volume (SV), and cardiac output (CO), that were combined to obtain biplane EDV, ESV, SV and CO values. The LA endocardium was traced at end systole in A2C and A4C views to calculate BP LA end systolic volume (LA Vol BP). Pulsed wave Doppler images taken at the mitral valve inflow tract in the A4C view were used to measure early (E) wave and late (A) wave peak filling velocities, E wave deceleration slope, E wave deceleration time, and the E/A ratio. Tissue Doppler imaging of the lateral annulus of the mitral valve in the A4C view was used to assess early (e’) and late (a’) mitral annular descent velocities, derive the e’/a’ ratio and complete the calculation for the E/e’ ratio, an index of LV filling pressure^44^. Pulsed wave Doppler images taken at the aortic outflow tract were used to measure aortic valve mean (AV Vmean), max (AV Vmax) and velocity time integral (AV VTI). The LV outflow tract diameter (LVOT) at beginning systole was measured using the parasternal long axis view. Body surface area (BSA, m^2^) at the time of echocardiogram was calculated as BSA = Body weight^2/3^ × 0.0969, utilizing the animal’s body weight (kg)^45^. In some animals, complete echocardiograms were not obtained due to difficulties in complete visualization of cardiac structures.

### Ex vivo: Immunostaining and Image Capture

Paraformaldehyde fixed, paraffin-embedded sections of longitudinally oriented LV (n=29), IVS (n=26), and RV (n=26) were serially sectioned and processed for histologic and immunohistochemistry (IHC). One section from each area was stained with Masson’s trichrome to assess fibrosis extent. Appropriate monoclonal antibodies were identified and protocols and procedures were developed and optimized for IHC staining of the phenotypes of interest. Subsequent serial sections were immunostained using a Leica Bond RX autostainer (Leica Biosystems, Inc, Buffalo Grove, IL) for CD31 (LV n=26, IVS n= 24, RV n=22) to assess endothelium/capillary density, (1:50, clone JC70A, Thermo Scientific, Waltham, MA), S100A4 (LV n=25, IVS n= 26, RV n=22) for fibroblasts and cells undergoing epithelial to mesenchymal transition (1:1000, clone CPR2761(2), Abcam, Cambridge, MA), CD68 (LV n=25, IVS n= 26, RV n=22) for macrophages (1:800, clone KP1, Dako/Agilent, Santa Clara, CA), and CD163 (LV n=26, IVS n= 25, RV n=22) for macrophages (1:800, clone EDHu-1, Thermo Scientific, Waltham, MA), all detected using the Vector Red alkaline phosphatase substrate detection kit (Vector Laboratories, Inc, Burlingame, CA). Finally, sections were dual stained with CD163 (Vector Red) and either pSTAT1 (DAB)(1:25, clone 58D6, Cell Signaling Technologies, Danvers, MA) or c-Maf (DAB)(1:50, clone EPR16484, Abcam, Cambridge, MA) to assess M1 or M2 macrophages as previously described.^46^ All slides were digitally scanned using a 20X objective on a BX61VS microscope and acquired on a VS 120 virtual slide imaging system (Olympus Corporation, Tokyo, Japan).

### Tissue Section Image Analysis

Acquired images were assessed using semi-automated algorithms developed internally specifically for phenotypes of interest on the Visiopharm Integrator System (Visiopharm, Broomfield, CO). First, a region of interest (ROI) was manually outlined around the myocardium, excluding valvular insertion points, the epicardium, papillary muscles and superficial large coronary arteries. The ROI was subsequently assessed via an automated algorithm that determined tissue area from non-tissue area (e.g. space around tissue or induced by artifactual shrinkage) using a green pixel filter thresholded to the background for each slide. The ROI was then quantitated by custom-designed algorithms specific to the stain or antibody of interest. For trichrome stains, the blue stained area representing fibrosis was quantified after defining the range of blue pixel values and was subsequently divided by the total tissue area previously obtained to calculate percent fibrotic area. For IHC stains for CD31, S100A4, CD68, and CD163, the positive area of interest was determined via thresholding using a red-green pixel contrast filter and divided by the total tissue area to calculate positive percentage area for CD31 or a normalized area (corrected for tissue area) for S100A4, CD68, and CD163. Additional steps to quantitate dual stained pSTAT1/CD163 and c-Maf/CD163 sections included quantifying the total CD163 positive area as described above, then running an algorithm that outlined CD163 stained areas to create new ROIs. The DAB positive area within these ROIs was thresholded using the DAB specific filter developed by Visiopharm. ROIs containing positive DAB staining were subsequently converted to a dual stain positive area and quantified. Finally, dual stained pSTAT1/CD163 and c-Maf/CD163 positive areas were normalized to the mean tissue area.

### Statistics

All analyses were performed using Statistica ver. 13.3 (TIBCO Software, Palo Alto, CA). All data were analyzed for normality and found to be normally distributed. Age, body weight, and BSA phenotypes were analyzed using Student *t* tests. Echocardiographic phenotypes were analyzed via analysis of co-variance (ANCOVA) and least square differences (LSD) post-hoc tests using BSA as a covariate. Histochemical and immunohistochemical outcomes were also covaried for BSA at time of death via ANCOVA and LSD post-hoc tests. Statistics are presented based on BSA corrected values, means ± SEM of the raw data not corrected for BSA are presented in the tables. The level of significance was set at p≤0.05.

## RESULTS

### Echocardiographic Phenotypes of Long Term Survivors

Structural and functional differences between LT IRR and LT CTL animals were assessed using B mode, M mode, and pulsed wave and tissue Doppler echocardiography (Supplementary Table 1). LT IRR animals differed from LT CTL in several diastolic function measures including increased transmitral early filling velocities (E) and E wave deceleration slope and decreased early mitral annular descent velocities (e’) that resulted in higher E/e’ ratios and decreased early (e’) to late (a’) mitral annular descent velocity ratios (e’/a’) (all p≤0.05) (Figure 1). The remaining diastolic function measures of transmitral late filling velocity (A), a’, and E/A ratio were not different between groups. Stroke volume (biplane and apical 4 chamber views), cardiac output (apical 4 chamber view), and end diastolic volume (apical 4 chamber views), were lower in LT IRR than LT CTLs (all p ≤ 0.05) and biplane measures of cardiac output and end diastolic volume tended to be lower as well (both p=0.07) (Figure 2). M-mode measures of cardiac structure corroborated our previous findings of decreased left ventricular diameters at end systole (LVIDs) and diastole (LVIDd) in LT IRR animals compared to LT CTLs, and also showed trends for decreased LV posterior wall thickness at end systole (LVPWs) (p=0.07) and diastole (LVPWd) (p=0.06) (Supplementary Table S1). No differences were observed in interventricular septal thicknesses at end systole (IVSs) or end diastole (IVSd) (both p> 0.5). Parasternal long axis measurement of the LV outflow tract diameter (LVOT) showed that LT IRR animals had significantly narrower outflow tract diameters compared to LT CTL (p=0.01) (Supplementary Table S1). No differences in heart rate, blood pressures (systolic, diastolic, and mean), ejection fraction, fractional shortening, or left atrium volume and size measures; aortic root to left atrium ratio (AO/LA) and left atrial end systolic volume (biplane) (LA Vol BP) were found.

**Figure 1.**
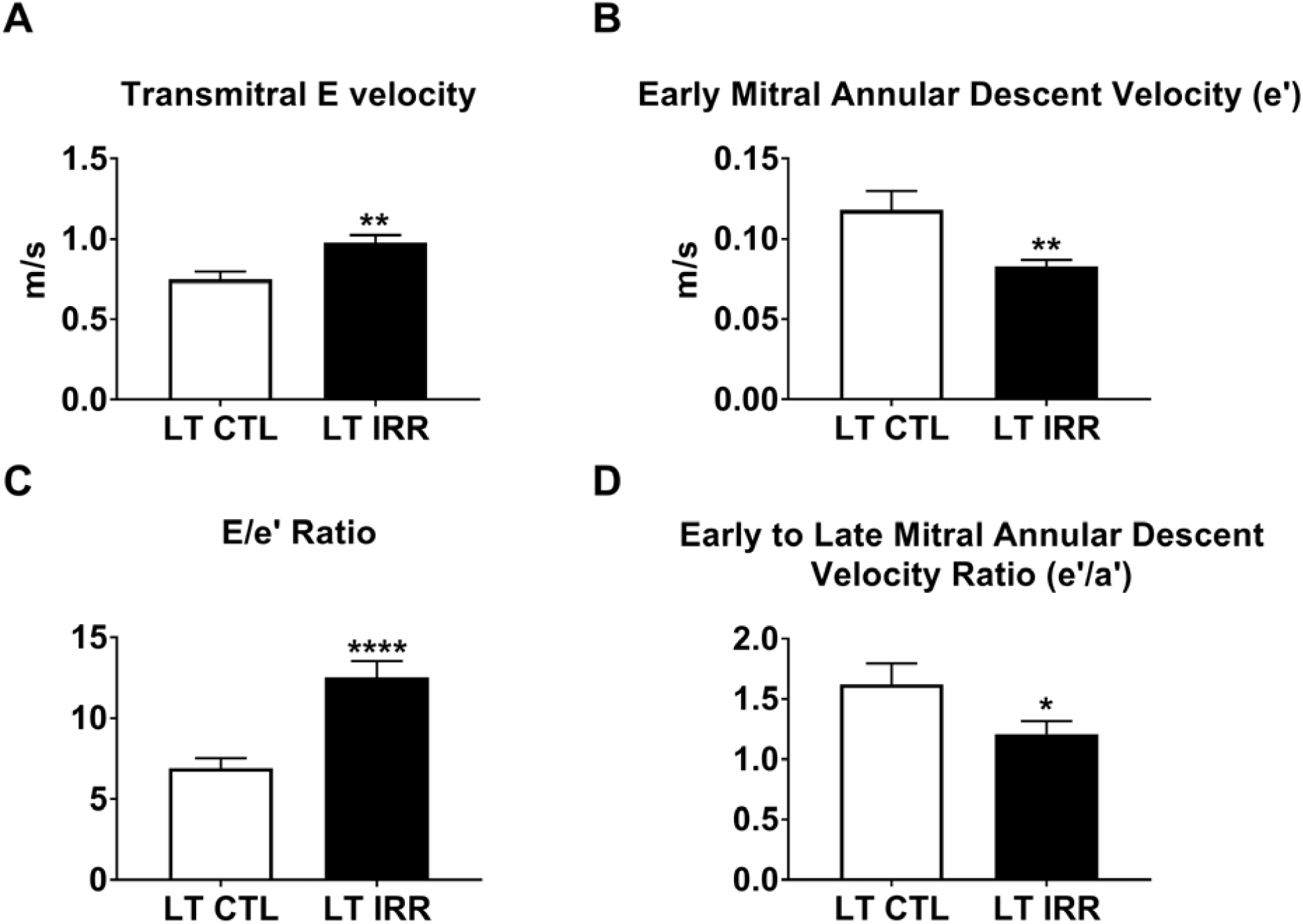
Echocardiographic Diastolic Function in Long Term Irradiated (LT IRR) and Long Term Controls (LT CTL): LT IRR animals had significantly increased transmitral early filling peak velocity (E) (Panel A), decreased early mitral annular descent velocity (e’)(Panel B), increased E/e’ ratio (Panel C) and decreased early to late mitral annular descent velocity ratio (e’/a’)(Panel D) compared to LT CTL animals. Data are mean ± SEM. *p<0.05, **p<0.01, ****p<0.0001

**Figure 2.**
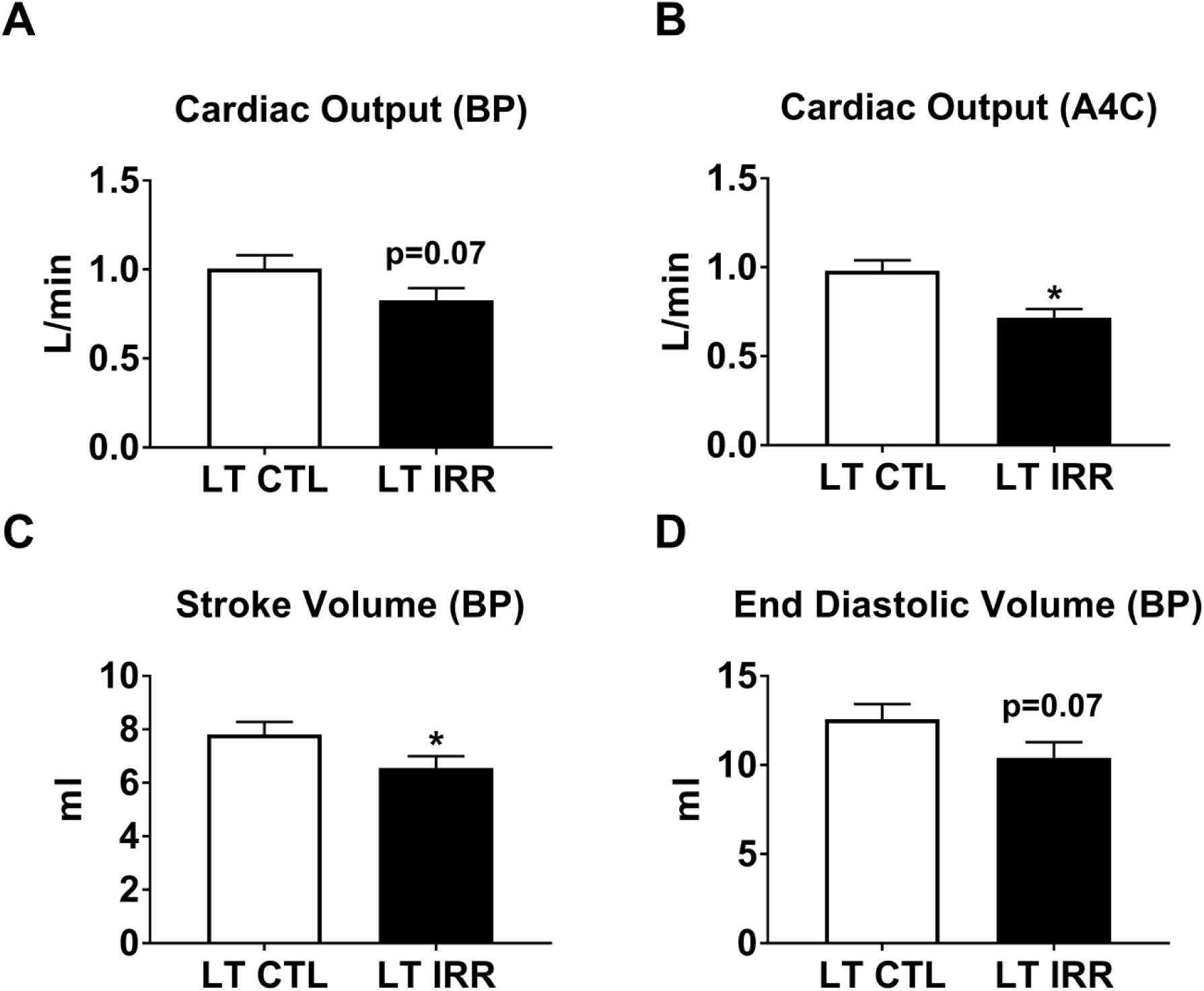
Echocardiographic Cardiac Volume Measures in Long Term Irradiated (LT IRR) and Long Term Controls (LT CTL): LT IRR animals tended to have decreased cardiac output calculated by method of disks measurements using the perpendicularly oriented apical 2 chamber (A2C) and apical 4 chamber (A4C) views resulting biplane measurement (BP) (Panel A). LT IRR animals did have significantly lower cardiac output when only the A4C chamber view was used (Panel B). Additionally, LT IRR animals had decreased stroke volume (BP measurement) and a tendency for decreased end diastolic volume (BP measurement). Data are mean ± SEM. *p<0.05

### Echocardiographic Phenotypes of Short Term Survivors

Structural and functional differences between ST IRR and ST CTL animals were evaluated similarly as LT IRR vs LT CTLs (Supplementary Table S2). Like LT IRR animals, ST IRR animals also had increased E/e’ ratios and decreased e’ and e’/a’ ratios compared to ST CTLs (all p≤0.05) (Figure 3), while E, A, E/A ratio, and a’ were not significantly different between groups. ST IRR and ST CTL did not differ in cardiac output, stroke volume, end diastolic volumes, heart rate, blood pressures (systolic diastolic, and mean), ejection fraction, fractional shortening, IVSd, ESV, LVIDs, LVIDd, AO/LA, or LA Vol BP. Interestingly, ST IRR animals had significantly thickened LVPWs, LVPWd, and IVSs (all p≤0.05) compared to ST CTL (Supplementary Figure 1).

**Figure 3.**
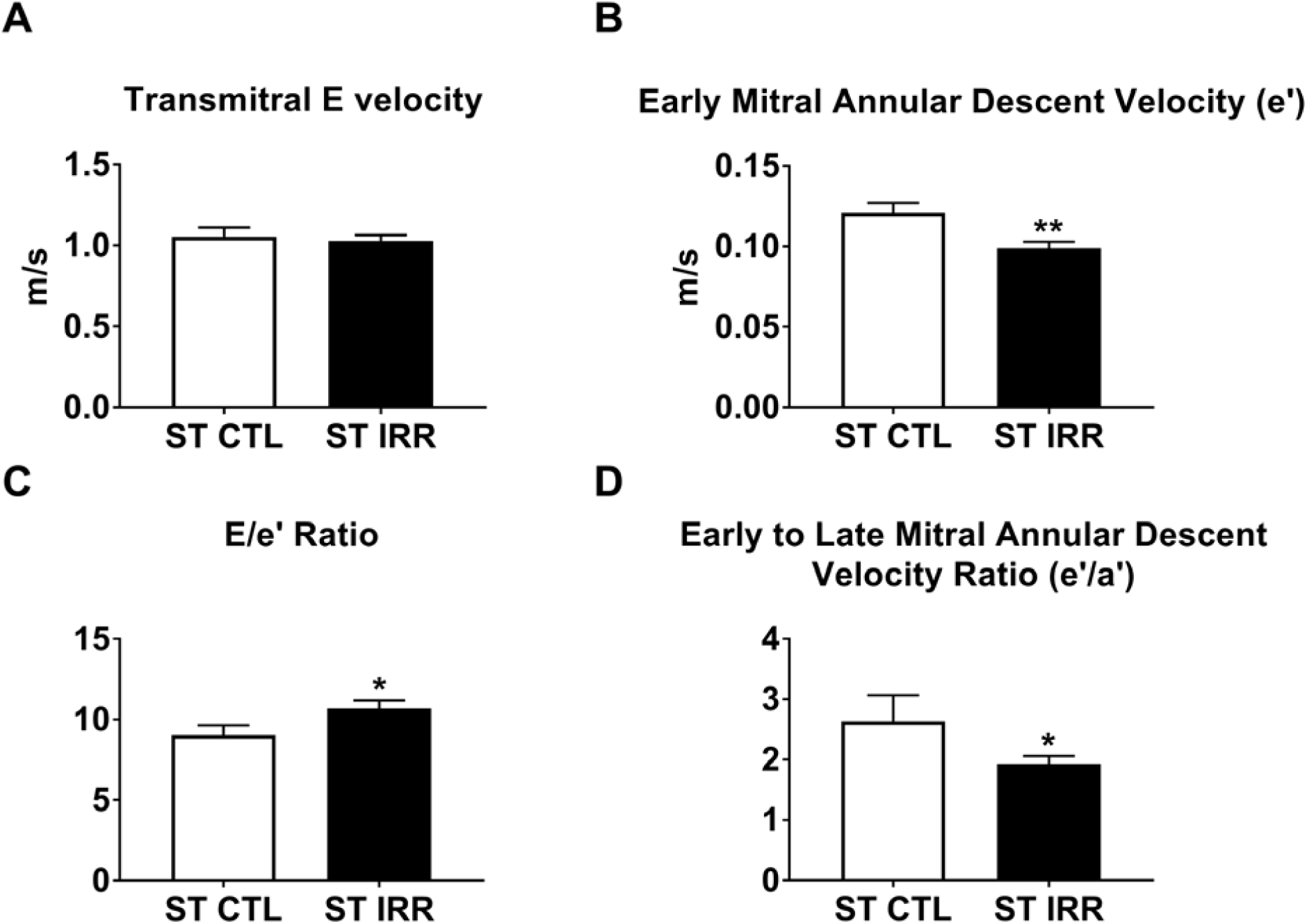
Echocardiographic Diastolic Function in Short Term Irradiated (ST IRR) and Short Term Controls (ST CTL): ST IRR animals had no differences in transmitral early filling peak velocity (E) (Panel A), but decreased early mitral annular descent velocity (e’)(Panel B), increased E/e’ ratio (Panel C) and decreased early to late mitral annular descent velocity ratio (e’/a’)(Panel D) compared to ST CTL animals. Data are mean ± SEM. *p<0.05, **p<0.01

### Histological and Immunohistochemical Staining of the Left Ventricular, Interventricular Septal, and Right Ventricular Myocardium

Sections of LV, IVS, and RV from 17 deceased previously irradiated animals (Deceased IRR) and 12 deceased controls (Deceased CTL) were evaluated for structural and cellular components using histochemical and immunohistochemical stains for fibrosis (Masson’s trichrome), capillary density (CD31), fibroblasts (S100A4), macrophages (CD68 and CD163) and M1 or M2 macrophages (pSTAT1/CD163 and c-Maf/CD163 respectively). IRR animals had significantly greater positive trichrome staining of the LV (p<0.05), and no significant difference in the RV or IVS compared to CTL animals. (Figure 4) Capillary density was reduced in the IRR group in the LV (p<0.05) and IVS (p<0.01), but not the RV. (Figure 4) IRR myocardium exhibited increased S100A4^+^ area in all cardiac sections (LV p<0.01, IVS p<0.001, and RV p<0.05) compared to CTL (Figure 5). In addition, approximately 100 fold less CD68^+^ macrophage staining area (LV mean= 8900 um^2^, IVS mean=6134 um^2^, RV =3484 um^2^) was observed relative to CD163^+^ macrophage staining area (LV mean= 905232 um^2^, IVS mean= 720474 um^2^, RV mean= 339053 um^2^) across all animals, and CD163 staining frequently co-localized with CD68 staining (Figure 6). Based on these findings we subsequently used CD163 as a pan-macrophage marker and used nuclear staining pSTAT1 or c-Maf to indicate M1 or M2 like macrophages respectively. Deceased IRR animals had increased CD163^+^ macrophage staining in the LV (p<0.05), IVS (p<0.01), and RV (p<0.05), increased c-Maf^+^/CD163^+^ macrophages in the LV (p<0.05) and IVS (p<0.01), but not RV, and increased pSTAT1^+^/CD163^+^ macrophages in the LV (p<0.01), IVS (p<0.01), and RV (p<0.05). (Figure 8) Finally, we compared the c-Maf^+^/CD163^+^ (M2) to pSTAT1^+^/CD163^+^ (M1) ratio and found that IRR animals had a significant decrease in the M2/M1 ratio across all assessed regions of the heart (all p<0.05).(Figure 7)

**Figure 4.**
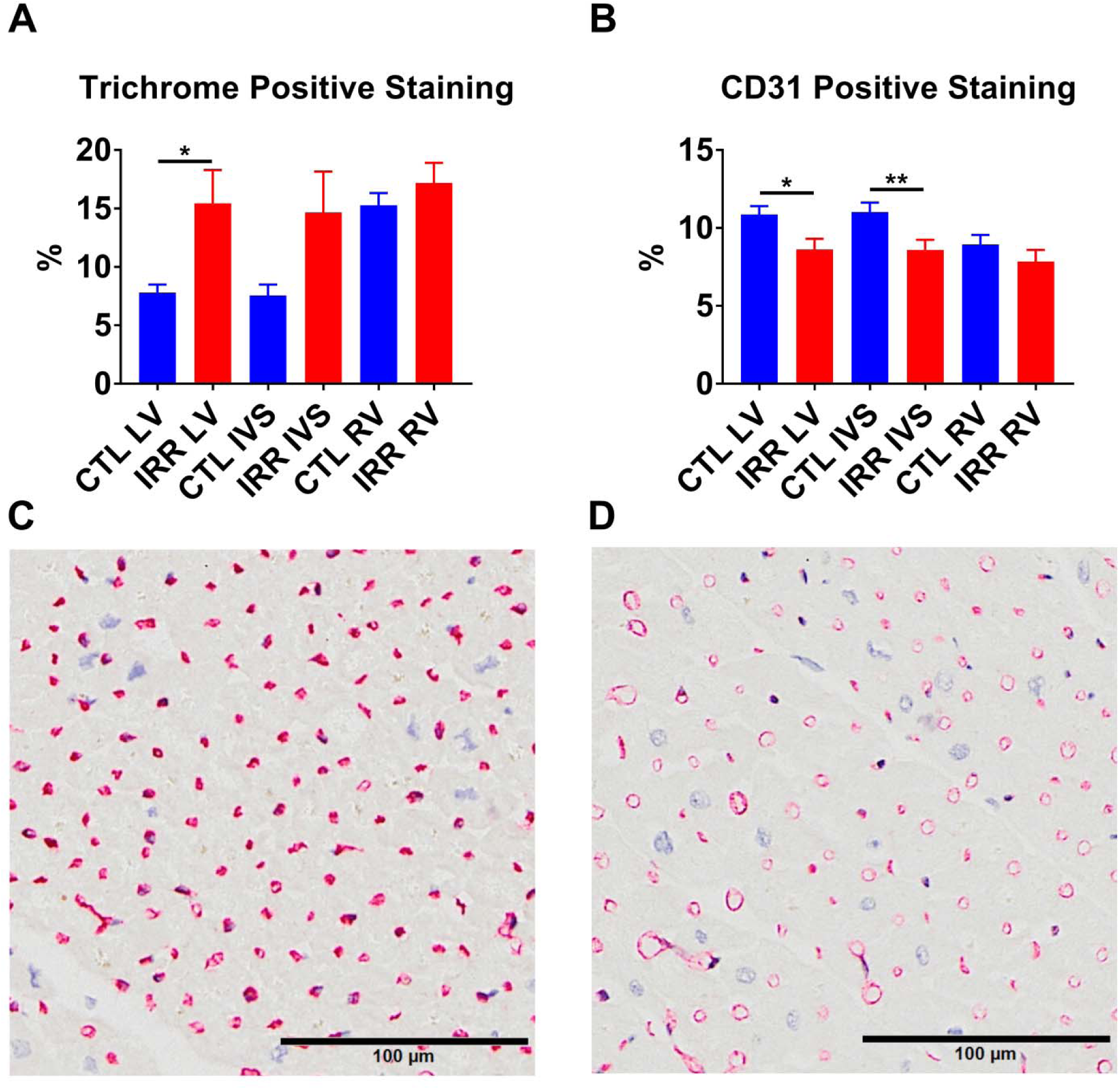
Fibrosis and Capillary Density Immunostaining in Deceased Irradiated (IRR) and Control (CTL) Animals: IRR animals had significantly increased percent positive trichrome staining of the left ventricle (LV) but not the interventricular septum (IVS) or right ventricle (RV) compared to CTL. IRR animals also had significantly decreased CD31 percent positive staining in the LV and IVS, but not RV compared to CTL (Panel B). Panels C and D are representative CD31 stained sections of LV mid-myocardium from a CTL and IRR animal respectively, demonstrating increased CD31 staining in CTLs. *p<0.05, **p<0.01

**Figure 5.**
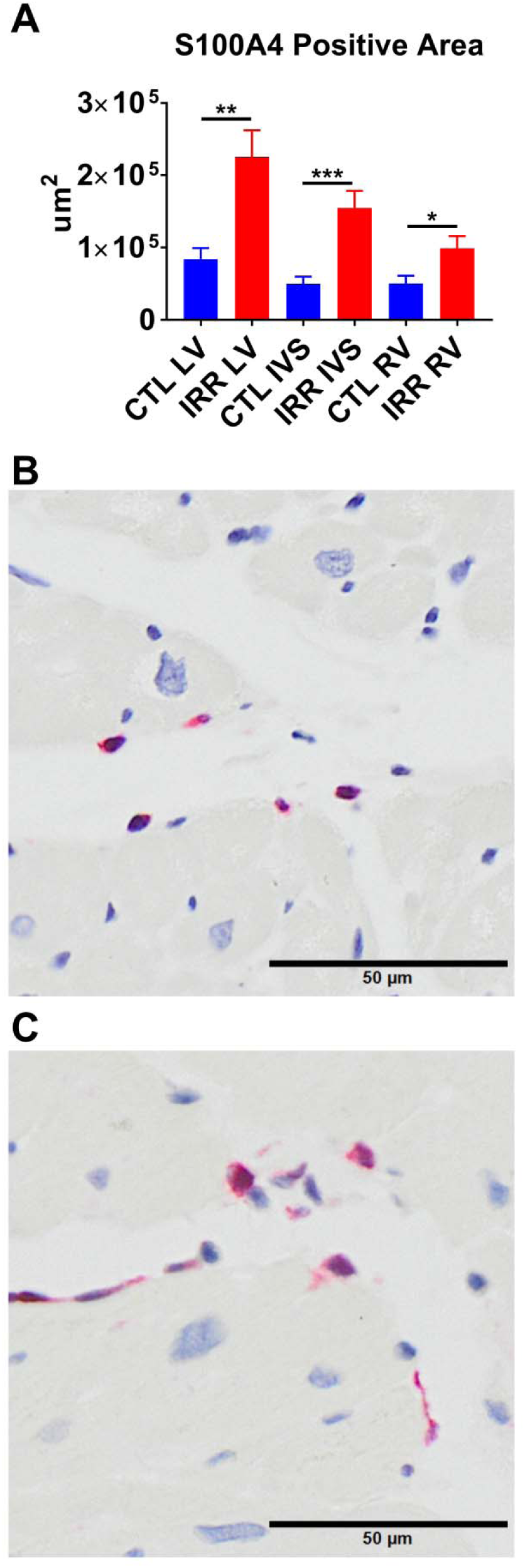
Fibroblast Immunostaining in Deceased Irradiated (IRR) and Control (CTL) Animals: IRR animals had greater S100A4 staining in all evaluated sections of myocardium compared to CTL animals (Panel A). Panels B and C are representative mid-myocardial S100A4 stained sections from a CTL and IRR animal respectively illustrating typical positive staining increases in IRR animals. *p<0.05, **p<0.01, ***p<0.001

**Figure 6.**
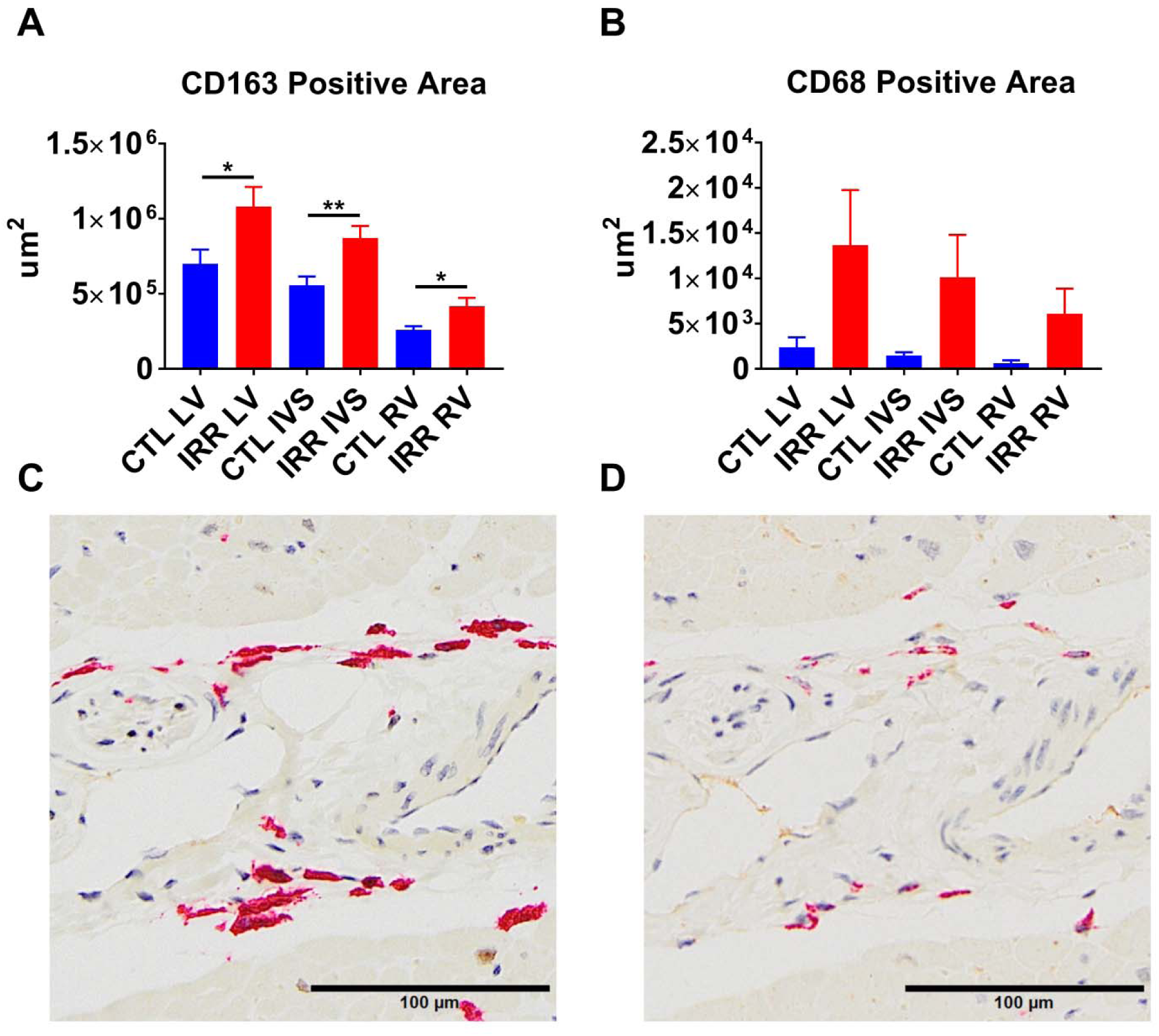
Macrophage Immunostaining in Deceased Irradiated (IRR) and Control (CTL) Animals: IRR animals had increased normalized CD163 positive staining area in the left ventricle (LV), interventricular septum (IVS), and right ventricle (RV) compared to controls (CTL) (Panel A). In contrast no differences were observed in CD68 staining which was highly variable across both groups (Panel B). Panels C and D are CD163 and CD68 stains respectively from the same animal in the same location illustrating that CD68^+^ macrophages were also often CD163^+^. *p<0.05, **p<0.01

**Figure 7.**
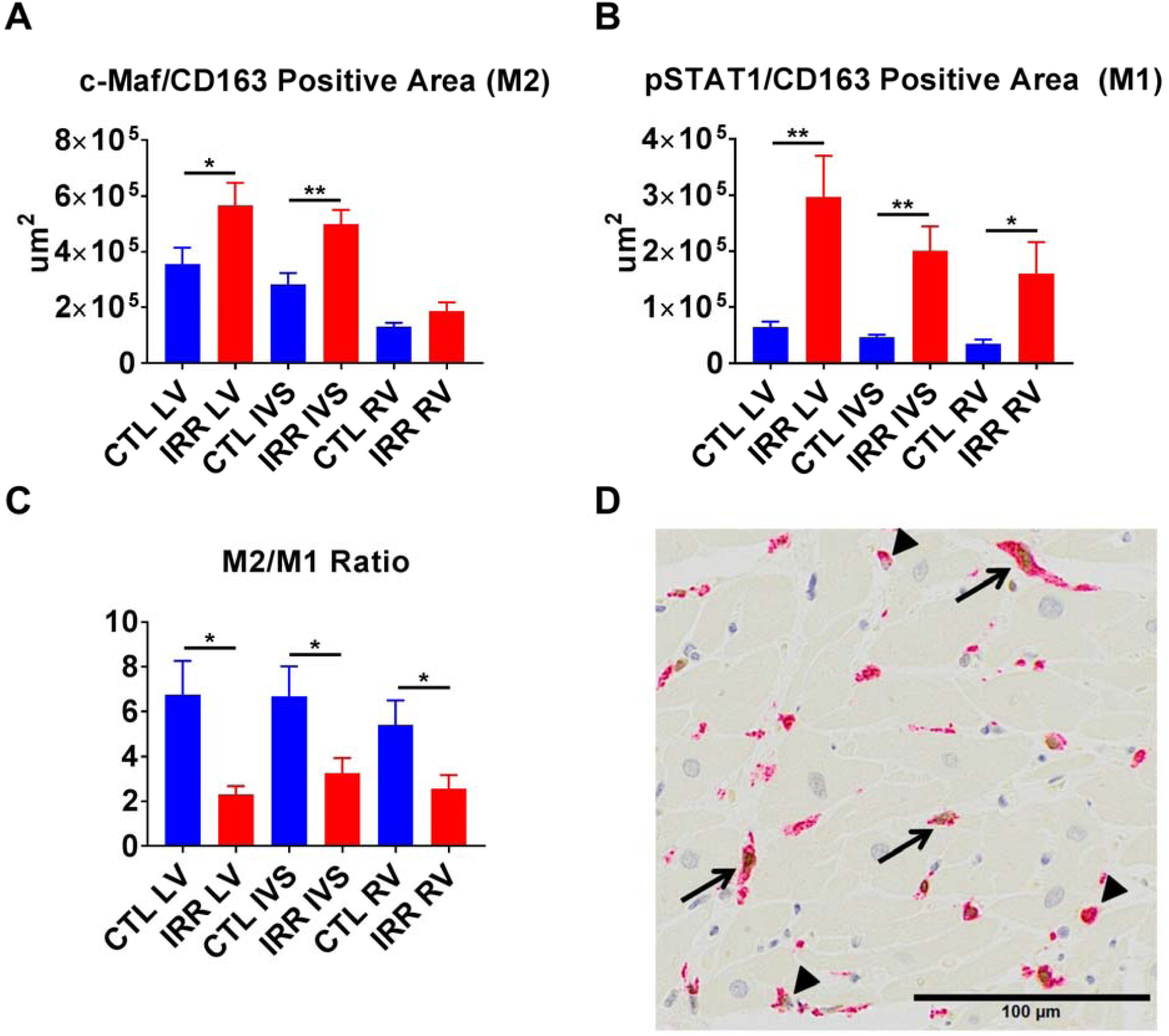
Macrophage Polarization in Irradiated (IRR) and Control (CTL) Myocardium: IRR animals had increased c-Maf^+^/CD163^+^ staining (M2 polarization) in the left ventricle (LV), interventricular septum (IVS), but not right ventricle (RV) compared to controls (CTL) (Panel A). Increased pSTAT1^+^/CD163^+^ staining (M1 polarization) was noted in all sections (LV, IVS, and RV) in IRR animals compared to CTL (Panel B). This translated to a decrease in the M2/M1 macrophage ratio across all sections in IRR animals (Panel C). Panel D is a representative dual stained image indicating dual stained c-Maf^+^/CD163^+^macrophages (arrows) and single stained CD163^+^ macrophages (arrowheads) from the LV of an irradiated animal. *p<0.05, **p<0.01

## DISCUSSION

The present study demonstrates delayed effects of acute ionizing radiation exposures on the heart including 1) diastolic dysfunction that was worse in long term than short term survivors, 2) decreased cardiac volumetric measures in long term survivors, 3) increased left ventricular fibrosis, 4) decreased left ventricular and interventricular septal capillary density, and 5) increased fibroblast, pan-macrophage, M1 macrophage, and M2 macrophage staining in all cardiac regions, with a greater relative increase in M1 macrophages. Our most consistent finding was left ventricular diastolic dysfunction (LVDD), which was present in both younger (4.7 to 6.0 years old) short-term survivors (ST IRR) (2.16 to 2.96 years post-irradiation) and worsened in older (12.6 to 20.3 years old) long-term survivors (LT IRR) (9.7 to 11.6 years post-irradiation) compared to age matched controls for each group. Both ST IRR and LT IRR groups had decreased early mitral annular descent velocities (e’), one of the earliest and consistent echocardiographically derived indicators of LVDD.^47^ This measure of LV relaxation decreases with increased myocardial stiffness and is generally not influenced by preload, unlike transmitral early filling peak velocity (E)^48^ which was elevated in LT IRR relative to LT CTL but did not differ between ST IRR and ST CTL. E can be corrected for the influence of preload and relaxation by dividing it by e’, and the resulting E/e’ ratio has been demonstrated to be the best index to detect LVDD (86%) in heart failure with preserved ejection fraction (HFpEF) in human patients who also had LVDD diagnosed via invasive conductance catheter derived pressure-volume loop (PVL) analyses.^49^

Our data suggests that diastolic function may decline fairly soon after TBI exposure, which has also been noted in human breast cancer patients receiving contemporary radiotherapy ^57^, and that effects on cardiac volumetric measures may require more time to develop as we only observed differences in the LT IRR vs. LT CTLs.

In addition to differences in diastolic function, we also found significant differences in echocardiographically determined cardiac structure. LT IRR animals had decreased LV internal diameters (LVIDd and LVIDs) compared to LT CTL, which corroborates our previous work in the RSC that used 13/20 of the LT IRR and 11/11 LT CTL animals reported here.^23^ In contrast, we did not detect differences in LV internal diameters in ST IRR compared to ST CTLs, which were younger and more recently irradiated, although there were significant differences between ST IRR and ST CTL in thickness of the LV posterior wall at end systole and diastole and the IVS at end diastole. Interestingly, these cardiac dimensions are nearly identical in LT IRR and ST IRR animals, whereas ST CTL animals have numerically smaller LV posterior wall and IVS thicknesses compared to LT CTLs, accounting for the findings of significance. This difference may be accounted for by a relatively young age of the ST cohort monkeys, as male rhesus macaques do not plateau in body size until around 10 years of age.^58^

Among deceased subjects, LV fibrosis extent was significantly increased in IRR compared to CTL animals and tended to be increased in the IVS, but not in the RV. However, the RV also had the highest percentage of fibrosis in both comparison groups, perhaps a function of the thinner RV wall compared to LV and IVS. The difference in fibrosis extent by myocardial region may also be related to reduction in capillary density, which has been demonstrated as a late effect in the LV of mice ^5, 6^, rats ^6^, and rabbits ^7^. Similar to the findings in these models, we observed a decrease in CD31 staining in the LV, and additionally the IVS, but not the RV.

Cardiac fibroblast content assessed using S100A4 (aka fibroblast specific protein 1, FSP1) was significantly elevated in all regions of TBI myocardium at a level of 10x that of CD68^+^, the most commonly referenced macrophage marker colocalizing with S100A4 ^60, 63^, indicating that the majority of S100A4 staining was due to cardiac fibroblast expansion.

Macrophage markers CD68 and CD163 were both elevated in myocardium from TBI subjects. CD163 appears to be a better general macrophage marker than CD68^46^, so we used CD163 as a pan-macrophage marker and used dual staining and co-localization IHC with pSTAT1, a transcription factor elicited by interferon signaling^66^, to specifically identify pro-inflammatory M1 polarized macrophages, or c-Maf, a transcription factor necessary for IL10 production^67^, to identify anti-inflammatory/reparative M2 polarized macrophages^46^. In both controls and TBI subjects, M2 staining phenotypes predominated over M1. TBI led to increased macrophage infiltration of the heart, with both M1 and M2 populations increased in comparable numbers, and a greater proportional increase in M1 macrophages due to a relatively sparse population of M1 macrophages in CTL myocardium.^46^ Collectively, this suggests that years after the initial IR exposure there is a long lived pro-inflammatory stimulus (M1) and repair (M2) by macrophage populations which leads to an accumulation of extracellular matrix and fibrosis.

Our studies are consistent with long term consequences of ionizing radiation on diastolic dysfunction recently tied to an increased risk of HF, specifically HFpEF,^56^ where women who had received contemporary radiotherapy for breast cancer (overall mean cardiac radiation dose of 3.3 ± 2.7 Gy, mean interval from radiotherapy to HF of 5.8 ± 3.4 years) had significantly increased odds ratios (OR) for HF (OR=9.14) and HFpEF (OR=16.9), but not HFrEF, even after adjustment for age, history of ischemic heart disease, history of atrial fibrillation, and cancer stage.^56^

Macrophages are master regulators of inflammation and fibrosis, as they play key roles in the initiation, production, and resolution of fibrosis.^64^ In the case of myocardial infarction, pro-inflammatory (M1) and anti-inflammatory/reparative monocyte/macrophages (M2) sequentially infiltrate murine hearts after infarction, M1 macrophages dominate the early phase before giving way to increasing M2 macrophages and extracellular matrix deposition.^65^ Depletion of circulating monocytes during the early stages after infarction impaired healing and increased necrotic debris accumulation, depletion during the later stages decreased collagen deposition and endothelial cells.^65^ We previously demonstrated that total body irradiation depletes circulating monocytes over the first month following irradiation which could lead to a similar outcome, with accumulated damage leading to a long term dysregulated inflammation and repair cycle which leads to fibrosis. Further studies are needed to explore this concept.

There are advantages and limitations inherent in our study design. Our nonhuman primate echocardiographic image acquisition and analyses were done in accordance with human echocardiography standards with minimal adaptation for rhesus macaques, allowing easy translatability of our findings. For the tissue analyses, quantification of immunohistochemical stains were unbiased, being tightly controlled based on background pixel values of acquired digital images and performed on the entire myocardium in each region. It is also worth noting that the non-irradiated control hearts available for histochemical and IHC analyses were derived largely from monkeys that were in declining health which led to their necropsy. In addition, macrophage polarization is more of a spectrum than a dichotomy, so our dual labeled immunohistochemical approach to identify “M1” and “M2” cannot fully characterize the macrophage phenotypes in the myocardium. Future studies using state of the art single nucleus RNAseq and other techniques will help to explore in depth the characteristics of the macrophages in the myocardium.

In summary, irradiated animals showed 1) diastolic dysfunction that was worse in long term than short term survivors 2), decreased cardiac volumetric measures in long term survivors 3), increased left ventricular fibrosis 4), decreased left ventricular and interventricular septal capillary density 5), increased fibroblast, pan-macrophage, M1 macrophage, and M2 macrophage staining in all cardiac regions, with a greater relative increase in M1 macrophages. These findings provide insight into the delayed effects of acute ionizing radiation exposure on the heart and indicate that novel therapies targeting the initiation, regulation, and resolution of myocardial inflammation and fibrosis by macrophages have potential to mitigate radiation induced heart disease and perhaps fibrosis in other tissues.

## ABBREVIATIONS

DEARE: Delayed effect of acute ionizing radiation exposure
RIHD: Radiation-induced heart disease
TBI: Total body Irradiation

## ACKNOWLEDGMENTS

We acknowledge the support of the Comparative Pathology Laboratory Shared Resources of the Comprehensive Cancer Center of Wake Forest Baptist Medical Center. We thank Jean Gardin, Cathy Mathis, Renae Hall, Russell O’Donnell, Chrystal Bragg, Matt Dwyer, Michael Bennett, J. D. Bottoms, and Kathy Stewart for their technical assistance.

---

**Figure S1.**
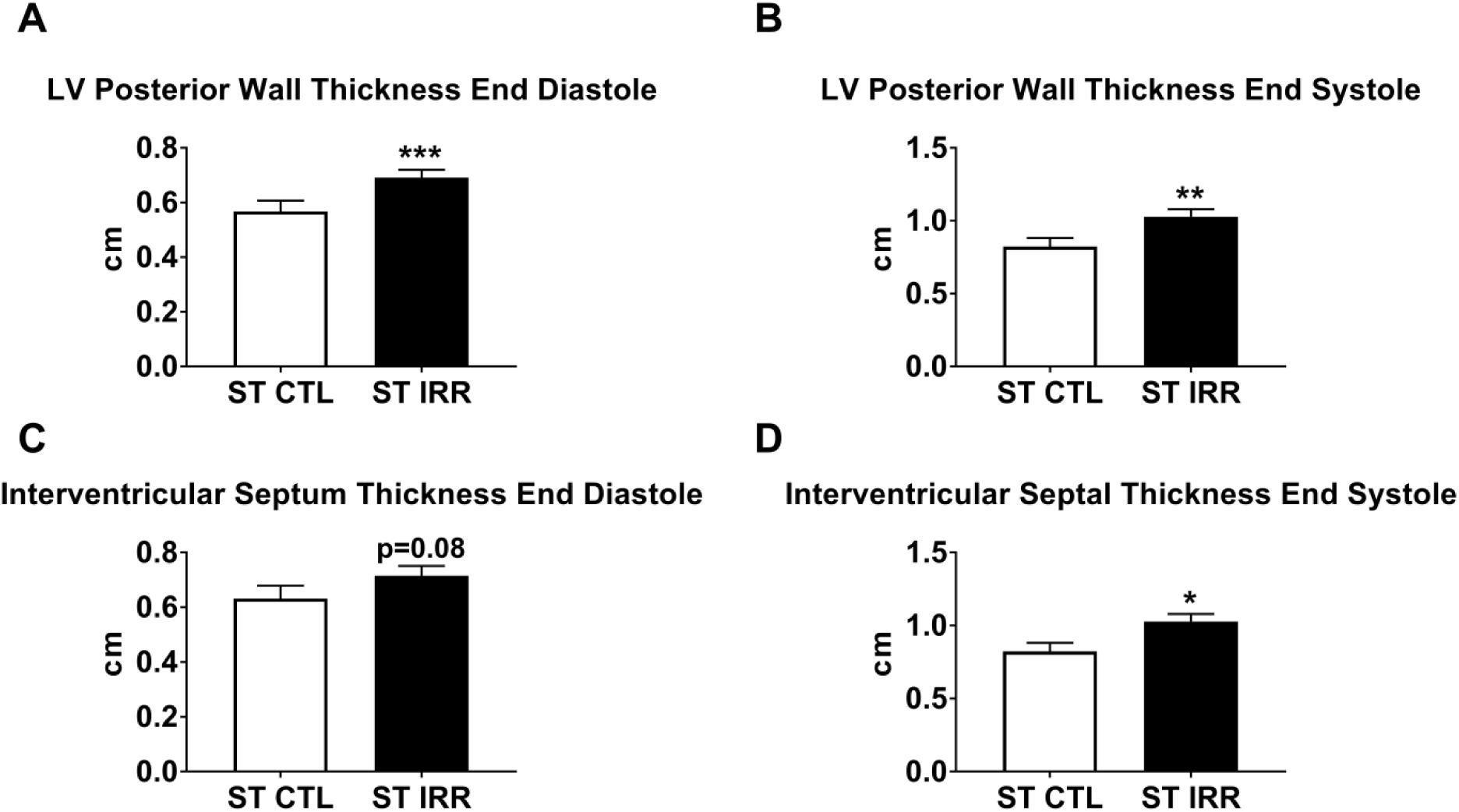
Echocardiographic Structural Difference in Short Term Irradiated (ST IRR) and Short Term Controls (ST CTL): ST IRR animals had an increased left ventricular posterior wall thickness at both end diastolic and end systole (Panels A & B), a tendency for increased interventricular septal thickness at end diastole (Panel C), and significantly increased interventricular thickness at end systole (Panel D). Data are mean ± SEM. *p<0.05, **p<0.01, ***p<0.001

